# Cochlear histopathology in macaques after noise-induced temporary threshold shifts

**DOI:** 10.1101/2025.11.14.688497

**Authors:** J. A. Mondul, C. A. Mackey, A. N. Conner, C. A. Alek, D. Pitchford, O. Rausis, L. Liberman, M. C. Liberman, R. Ramachandran, T. A. Hackett

## Abstract

Noise exposures causing transient hearing loss were previously considered benign. However, recent work has revealed that temporary noise-induced threshold shifts may be associated with long-lasting cochlear histopathology. One such effect is cochlear synaptopathy, i.e. changes to the afferent synapse between inner hair cells and auditory nerve fibers. Noise-induced synaptopathy has been extensively characterized in several rodent models, and temporal bone studies suggest similar age-related changes in humans. However, it remains unclear how noise-induced temporary threshold shifts affect cochlear structures in humans and nonhuman primates, which show greater resistance to noise exposure than other animals. Additionally, the long-term sequelae of temporary threshold shifts are largely unknown. Here, we characterized the effects of a noise exposure causing temporary hearing loss on cochlear histopathology in macaque monkeys at long post-exposure survival times. Overall, cochlear histopathology was variable across subjects, similar to the variable susceptibility observed in humans. At 2 and 10 months post-exposure, macaques had no significant loss of hair cells, inner hair cell synapses, or cholinergic efferent innervation. However, enlargement of ribbons in both inner and outer hair cells was observed. Together, these findings provide insight into the cochlear effects of single-exposure temporary threshold shifts in nonhuman primates.

**HIGHLIGHTS:**

- Macaques exposed to 120 dB SPL noise for 4h showed temporary threshold shifts
- Cochlear histopathology was evaluated at 2 and 10 months post-exposure
- Macaques had no significant loss of hair cells or inner hair cell synapses
- Chronic enlargement of inner and outer hair cell ribbons was observed
- Transient loss of outer hair cell ribbons was also observed

## 1. INTRODUCTION

High-intensity sounds induce large displacements of the cochlear sensory epithelium, causing mechanical and metabolic stress on the inner and outer hair cells (IHCs, OHCs). OHCs are generally more vulnerable than IHCs, and OHC loss is associated with permanent threshold shifts (PTSs) (Burton et al., 2020; Hamernik et al., 1989; Hawkins et al., 1976; Ryan & Dallos, 1975; Stebbins et al., 1979). However, PTS can also be seen with minimal loss of either IHCs or OHCs (Liberman & Dodds, 1984). Permanent damage to the stereocilia on surviving hair cells is a key contributor to this noise-induced threshold elevation (Engstrom, 1984; Liberman & Dodds, 1984; Wang et al., 2002).

In addition to hair cell loss, noise exposure can cause excitotoxic swelling of auditory nerve terminals (Le Prell et al., 2004; Puel et al., 1998; Robertson, 1983) and damage to or loss of presynaptic IHC ribbons, a pathology known as cochlear synaptopathy (SYN) (e.g., Furman et al., 2013; Kujawa & Liberman, 2009; Valero et al., 2017). SYN can also accompany OHC loss and permanent threshold shifts after acoustic overexposure (Fernandez et al., 2020; Valero et al., 2017) and can also occur without hair cell loss after exposures causing only temporary threshold shifts (Kujawa & Liberman, 2009; Lin et al., 2011). SYN is also seen in age-related hearing loss, where, at least in a mouse model, it appears before there are age-related threshold shifts or loss of sensory cells (Sergeyenko et al., 2013).

SYN reduces the number of auditory-nerve fibers innervating each IHC. Deafferentation does not elevate thresholds until it becomes severe (Makary et al., 2011; Schuknecht & Woellner, 1955; Wong et al., 2019), but it may underlie variability in hearing abilities among individuals with the same audiometric loss and could explain why some individuals with normal hearing sensitivity report hearing difficulties in background noise (i.e. hidden hearing loss; Bharadwaj et al., 2015; Cooper & Gates, 1991; Parthasarathy et al., 2020; Plack et al., 2016; Ruggles et al., 2011; Schaette & McAlpine, 2011).

Nonhuman primates are a useful model for human inner ear pathologies due to their phylogenetic proximity and comparable auditory anatomy and physiology. Like humans, nonhuman primates are less susceptible to acoustic injury than other commonly used animal models such as guinea pigs, chinchillas or mice (Burton et al., 2019; Valero et al., 2017). We previously demonstrated that macaque monkeys exhibit SYN after noise exposures causing large PTSs and massive OHC losses (Burton et al., 2020; Hauser et al., 2018; Mackey et al., 2021; Valero et al., 2017). Here, we investigated chronic cochlear histopathology following severe noise-induced temporary threshold shifts. We quantified OHC and IHC counts, OHC and IHC ribbon counts and sizes, and cholinergic olivocochlear efferent innervation density at 2 months and 10 months post-exposure to assess time-dependent effects of acoustic injury and recovery. This work provides critical insights into the dynamics and long-term progression of noise-induced inner ear damage in a nonhuman primate model.

## 2. MATERIALS AND METHODS

### 2.1 Subjects

Cochlear tissue was obtained from twenty-five young adult rhesus macaques (*Macaca mulatta*, 6-10 years old, 5 female). Subjects were randomly allocated to three different groups: unexposed controls (*n* = 10, 1 female), short-term post-exposure survival (2 months; *n* = 5), and long-term post-exposure survival (10 months; *n* = 10, 5 female).

Animals were maintained on a 12:12-h light:dark cycle. Veterinary assessments and experimental procedures occurred between 8:00 am – 5:00 pm during their light cycle. Four animals were socially housed. All other animals were individually housed due to incompatibility for social housing, but had visual, auditory, and olfactory contact with conspecifics maintained within the housing room. A commercial primate diet (Lab Diet 5037 or 5050, PMI Nutrition International, Brentwood, MO) was provided twice daily and was supplemented with fresh produce and/or foraging items (seeds, dried fruit, nuts, etc.). Animals were provided manipulanda as well as auditory, visual, and olfactory enrichment on a rotational basis. Filtered municipal water was provided at least once a day, as the animals were maintained on fluid restriction for study purposes. All animals were under the continuous care of veterinary staff and received semiannual comprehensive physical exams, including standard blood work (annual) and tuberculosis testing. Cranial implants (used for head fixation during psychophysical testing of parallel studies) were regularly cleaned with topical agents.

All research procedures were approved by the Institutional Animal Care and Use Committee at Vanderbilt University Medical Center.

### 2.2 Noise exposure and auditory physiological characterization

#### 2.2.1 Anesthetic procedures

Animals were anesthetized for auditory physiological testing and the noise exposure procedure. Initial sedation was induced with an intramuscular injection of ketamine (10 mg/kg) and midazolam (0.05 mg/kg), along with atropine (0.04 mg/kg) to minimize mucous secretions. Animals were intubated and anesthesia was maintained with isoflurane (1-2%). All anesthetized procedures were conducted in a sound-treated booth (Industrial Acoustics Corp, NY; Acoustic Systems, Austin, TX). Subjects were monitored intensively for a minimum of 72 hours post-procedure. Physiological testing was conducted 1-3 months prior to noise exposure, immediately after noise exposure (DPOAEs only, same session), 2 months post-exposure, and 9-10 months post-exposure.

#### 2.2.1 Noise exposures

Subjects underwent a single noise exposure intended to cause temporary threshold shifts. The noise exposure procedure was similar to that previously reported by our laboratory (Burton et al., 2020; Hauser et al., 2018; Mackey et al., 2021; Valero et al., 2017). The subject was laid prone on a table with the head slightly elevated in a sound treated booth. Closed-field speakers (MF1, Tucker-Davis Technologies) were coupled to the ears using 1.5” PE tubing and pediatric ER-3A insert earphones that were deeply inserted into each ear canal. Octave-band noise (2-4 kHz) was presented simultaneously to both ears at 120 dB SPL for four hours. The level of the exposure stimulus was calibrated using a 0.25” microphone (model 378C01, PCB Piezotronics Inc., Depew, NY) coupled to a 0.5cc coupler, and verified in the ear canal using a probe microphone system (Fonix 8000, Frye). Noise level varied by less than 0.3 dB SPL over the course of the four-hour procedure.

#### 2.2.2 Auditory brainstem response (ABR) testing

ABR recording methods were similar to those used in previous publications from our laboratory (Hauser et al., 2018; Stahl et al., 2022; Valero et al., 2017). Briefly, ABRs were measured using subdermal needle electrodes (Rhythmlink) placed on the mastoid (active), vertex (reference), and shoulder (ground) connected to a Medusa 4Z preamplifier (Tucker-Davis Technologies). Impedances for subdermal needle electrodes were consistently less than 1 kΩ. A closed-field speaker (MF1, Tucker-Davis Technologies) was coupled to the ear with a pediatric ER-3A foam tip.

Stimuli were created in SigGenRZ and generated by an RZ6 Multi-I/O Processor (Tucker-Davis Technologies). Stimuli (clicks, tonebursts from 0.5-32kHz) were presented at 27.7/s from 90-10 dB SPL with an alternating stimulus polarity for two separate runs of 1024 repetitions. Stimuli were calibrated (+/- 1 dB) using the same 0.25” PCB microphone and 0.5 cc coupler and verified in-ear using the Fonix probe microphone system. Stimulus presentation, signal acquisition, and data analysis was completed using BioSigRZ software. During online recording, the incoming signal was digitally filtered from 300-3000 Hz. During offline analysis, the two artifact-free waveforms were averaged, inverted, and low-pass filtered at 1500 Hz to product a single ABR trace per condition (2048 repetitions).

ABR threshold was defined as the lowest sound level that elicited a visually identifiable response greater than the noise floor (±20 nV). Peak-to-trough amplitudes for ABR Waves I-IV were visually identified for each trace to derive input-output functions.

#### 2.2.3 Otoacoustic emissions (OAE) testing

OAE testing was completed using a Scout Bio-logic OAE System (Natus, Pleasanton, CA). Recording methods were similar to those used in prior publications from our laboratory (Hauser et al., 2018; Stahl et al., 2022; Valero et al., 2017). A probe containing two speakers and one microphone was coupled to the ear with a pediatric foam tip. Distortion product otoacoustic emissions (DPOAEs; 2*f*_1_-*f*_2_) were measured in response to tone pairs (*f_2_* = 0.5-10kHz, 8 points per octave; *f_2_*/*f_1_* = 1.22; *L_1_*-*L_2_* = 10 dB; *L_1_* = 70-25 dB in 5 dB steps). As previously described (Stahl et al., 2022), DPOAE amplitudes and input-output functions were derived. DPOAE threshold was defined as the lowest sound level that elicited a DP amplitude > 0 dB SPL, and with a signal-to-noise ratio > 6 dB).

### 2.3 Cochlear histological preparations and quantification

Following completion of the physiological study, subjects were euthanized via overdose of sodium pentobarbital and sodium phenytoin (Euthasol; >120 mg/kg IV) and transcardially perfused with 0.9% phosphate-buffered saline and 4% phosphate-buffered paraformaldehyde (PFA). Temporal bones were extracted to harvest the cochlear tissue. The round and oval windows were opened, cochleas were perfused through the scala tympani with PFA, submerged in PFA for 2 hours, and transferred to 0.12 M EDTA for decalcification. Decalcified cochleas underwent dissection, imaging, and immunohistochemical analysis as previously described (Burton et al., 2020; Valero et al., 2017). Briefly, the organ of Corti and attached osseous spiral lamina was dissected into 10 – 11 pieces. Immunohistochemistry was used to label i) presynaptic ribbons (mouse IgG1 anti-CtBP2 (C-terminal binding protein 2); BD Transduction Labs; 1:200); ii) glutamate receptor patches (mouse IgG2 anti-GluA2; Millipore; 1:200), and iii) hair cell cytoplasm (rabbit anti-myo7a (myosin VIIa); Proteus Biosciences; 1:200). A fourth channel was used to label either cochlear afferent and efferent fibers (chicken anti-NFH (neurofilament-H); Chemicon; 1:1000) or cochlear efferent fibers (goat anti-choline acetyltransferase (ChAT); Millipore #AB144P; 1:100). Tissue was incubated in species-appropriate fluorescent secondary antibody conjugates (AlexaFluor), slide-mounted in Vectashield and coverslipped for confocal microscopy.

For synapse counts, the tissue was imaged on a Leica SP8 confocal microscope, using a 63X glycerol objective (1.3 N.A.), to acquire 3-dimensional image stacks at octave-spaced positions along the cochlear spiral from 0.125 to 32 kHz, with additional half-octave spacing in regions near the noise exposure band. The frequency correlate of each image stack was computed from a cochlear frequency map based on a Greenwood function (Greenwood, 1990), assuming an upper frequency limit of 45 kHz (based on Pfingst et al., 1978). Amira software (version 2019.4, Visage Imaging) was used to quantify IHC and OHC ribbons from confocal z-stacks using the *connected components* function, as described in prior publications (Valero et al., 2017). In the IHC area, orphan ribbons were distinguished from ribbon synapses using custom software that created a thumbnail array of the voxel space 1 micron around each ribbon, ordered by average pixel intensity in the GluA2 channel, followed by manual counting (Liberman et al., 2011). Synapse and ribbon counts in each z-stack were normalized to the number of hair cells, assessed by counting the number of nuclei (including fractions at the edges of the stack).

Hair cell survival was assessed in separate confocal z-stacks by counting cuticular plates normalized to the expected number of hair cells within each row. Amira software was used to quantify efferent terminal density underneath the OHCs (medial olivocochlear area) and IHCs (lateral olivocochlear area). Tunnel fibers were cropped out, and maximum intensity projection (MIP) images (XY plane) of the MOCs and LOCs were separately exported. Images were auto-thresholded in ImageJ to binarize the pixel intensities before quantifying the innervation area.

### 2.4 Statistical analyses

Statistical analyses were conducted using linear mixed effects models (“fitlme” in MATLAB 2018a). The dependent variable in the models was hair cell count, ribbon count, ribbon size, or efferent terminal density. Frequency and group (i.e., noise exposure status and post-exposure survival time) were entered as fixed effects into the model, while intercepts for individual subjects and ears were entered as random effects. In all cases, *p*-values were obtained by likelihood ratio testing of the model with the effect in question against the model without that effect. A significant *p*-value was defined as *p* < 0.05. *t*-statistics are reported for each model, similar to the *F*-statistic that is often reported for such models. Additional models that treated frequency bins (0.5-1.4 kHz, 2-5.6 kHz, 8-32 kHz) as a categorical variable were tested, but did not reveal any patterns different from the reported trends.

Ribbon volume distributions were compared using Kolmogorov-Smirnov two sample tests (“kstest2” in MATLAB 2018a). A Benjamini-Hochberg correction was applied to adjust the significance level for multiple comparisons (Benjamini & Hochberg, 1995). Estimates of ribbon volume range for a given group and frequency region were conducted using a jack-knife procedure. Data were resampled with a randomly selected 10% of the data left out. The number of resamples was set to equal the length of the data, approximately 5,000 on average.

## 3. RESULTS

### 3.1 Effects of noise exposure on cochlear function

Young-adult rhesus macaques were exposed to a 120 dB SPL octave band noise (2-4 kHz) for 4 hours while under anesthesia. This noise exposure was designed to cause severe temporary threshold shifts and cochlear synaptopathy, based on previous studies of noise-induced permanent thresholds shifts in rhesus macaques in our laboratory (Valero et al., 2017). Cochlear function was assessed before and after noise exposure using ABRs and DPOAEs (for comprehensive characterization, see Conner et al., 2025). Immediately after noise exposure, macaques had significant DPOAE threshold elevations across nearly all frequencies tested compared to pre-exposure measurements (Figure 1A, red filled vs. black symbols; post hoc *p* values < 0.05 for 0.56-9.1 kHz). By 2 and 9-10 months post-exposure, DPOAE thresholds were no longer different from pre-exposure values, except that thresholds were significantly lower (better) from 0.5-2 kHz (Figure 1A; post hoc *p* values < 0.05 for 0.5-1.9kHz at both time points). At 2 months and 9-10 months post-exposure, ABR thresholds were not significantly different from pre-exposure values (Figure 1B; *p* > 0.05). Interestingly, suprathreshold click ABR Wave I amplitudes were significantly larger (Figure 1C) at both 2 months (post hoc *p* values < 0.05 for 60-90 dB SPL) and 9-10 months post-exposure (post hoc *p* values < 0.05 for 70-90 dB SPL). Overall, these physiological measures demonstrated temporary noise-induced threshold shifts that recovered to normal, which was sustained through long post-exposure time points.

**Figure 1.**
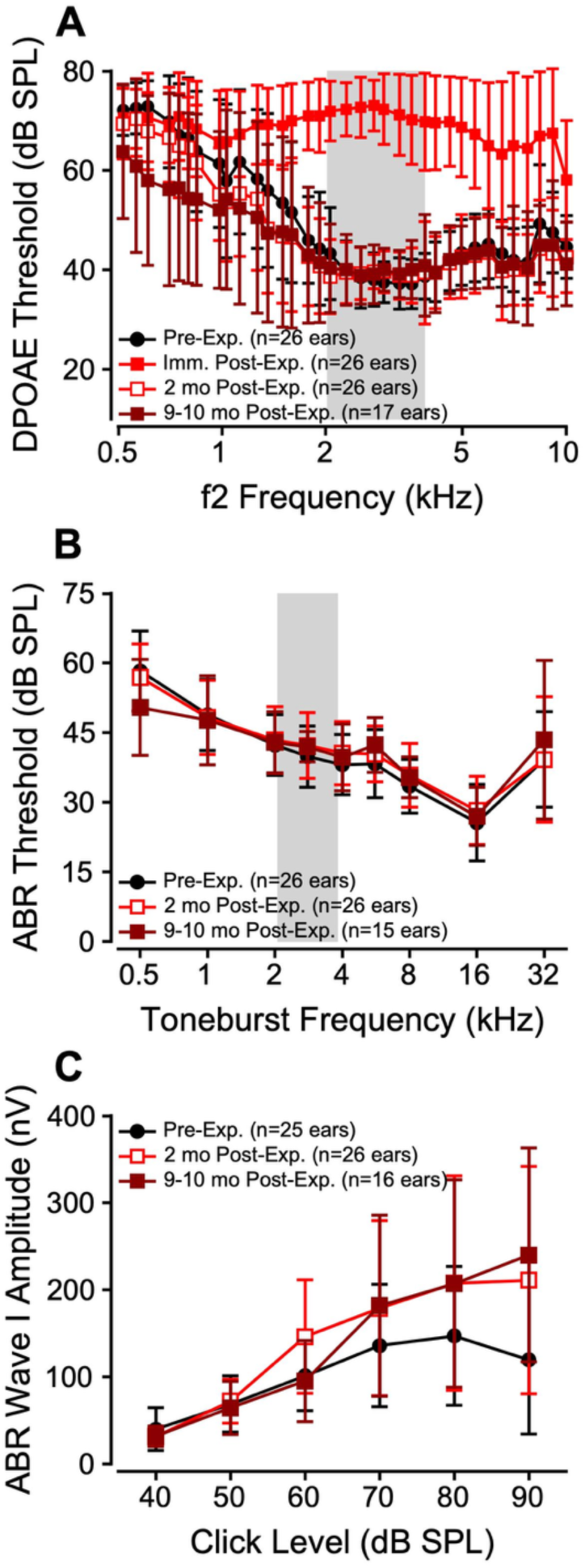
Physiological measures of auditory function in macaques before and after noise exposure. **A**: DPOAE thresholds as a function of f_2_ frequency. **B**: ABR thresholds as a function of toneburst frequency. **C:** ABR Wave I amplitude as a function of click level. Black circles = pre-exposure, filled red squares = immediately post-exposure, open red squares = 2 months post, dark red squares = 9-10 months post-exposure. Error bars indicate ± 1 standard deviation from the mean. Gray boxes illustrate the noise exposure band.

### 3.2 Noise-induced temporary threshold shifts caused minimal outer and inner hair cell loss

As seen in humans, macaques have three to four rows of OHCs and one row of IHCs (Figure 2; Bredberg, 1968; Johnsson & Hawkins, 1967; Lonsbury-Martin et al., 1988). Control ears had nearly full complements of OHCs and IHCs across the cochlea (Figure 2A; 2D-E, black symbols). Although the data suggest there might be a subtle loss of OHCs, especially in the long-surviving exposure group, the differences from control were not statistically significant. (Figure 2B-C; 2D, red open and dark red symbols; *t*(*df*) = -1.04(522), *p* = 0.30). Inter-subject and across-ear variability was substantial for OHC survival post-exposure, with 1 out of 10 short-term ears and 6 out of 20 long-term ears showing >10% OHC loss (Figure 2D, inset panel). The modest reduction (11% on average) in OHC survival for frequencies near the noise exposure band did not significantly affect DPOAE or ABR thresholds (Figure 1A-B). Little to no IHC loss was observed at 2 or 10 months post-exposure (Figure 2B-C; 2E; *t*(*df*) = 0.30(500), *p* = 0.76), although there was a significant effect of frequency driven by the 5-10% IHC loss at 32kHz across all groups (*t*(*df*) = -9.38(500), *p* = <0.001).

**Figure 2.**
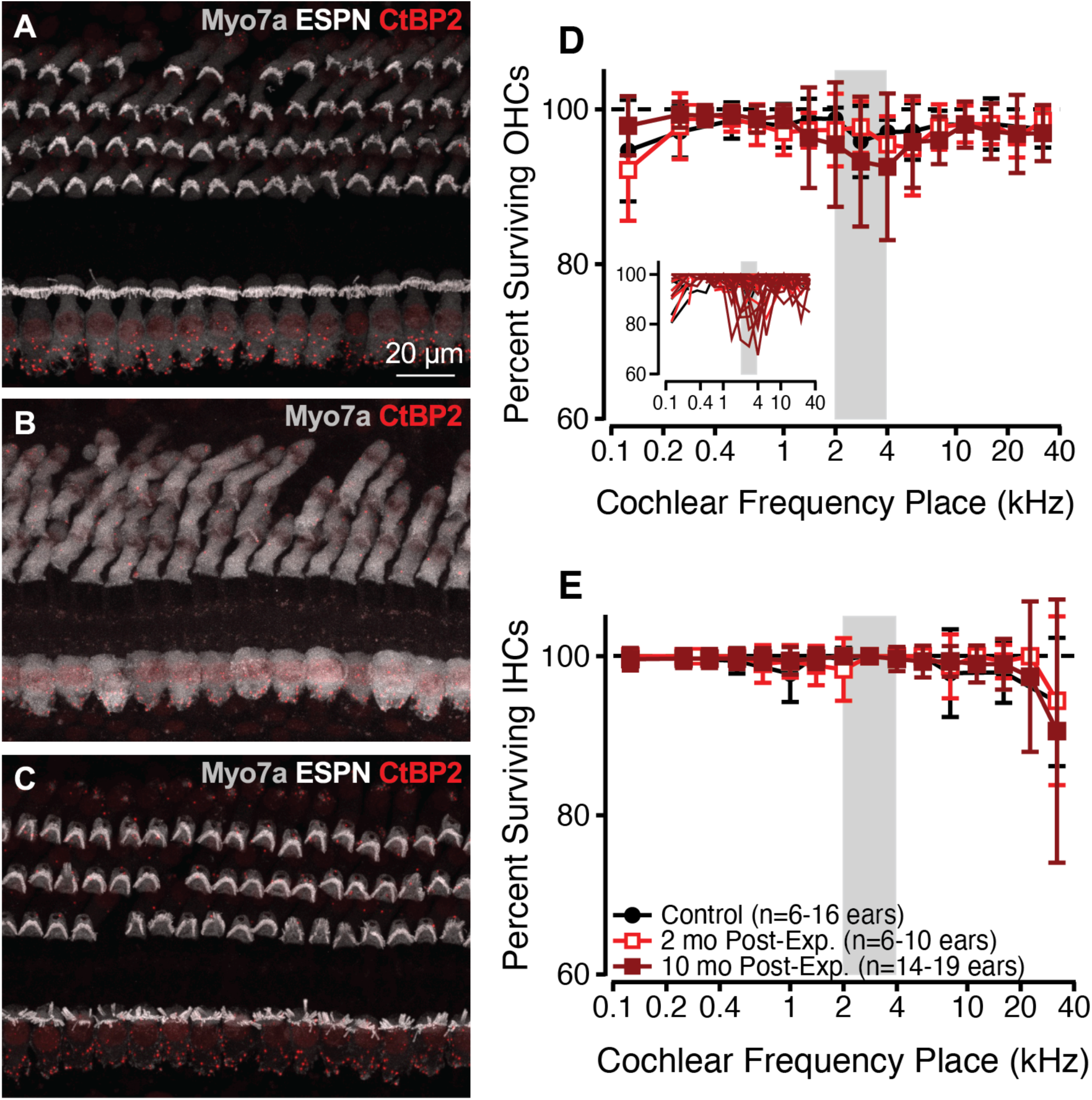
**A-C:** Confocal maximum intensity projection images of outer and inner hair cells at the 5.6 kHz place in a control (**A**) and in noise-exposed subjects at 2 months (**B**) and 10 months (**C**) post-exposure. Outer hair cell survival (**D)** and inner hair cell survival (**E**) as a function of cochlear frequency place in controls (black circles, *n* = 6-16 ears per frequency) and at 2 months (open red squares, *n* = 6-10 ears) and 10 months (dark red squares, *n* = 14-19 ears) post-exposure. Inset on panel D illustrates variability among individual ears. Error bars indicate ± 1 standard deviation from the mean. Gray boxes illustrate the noise exposure band.

### 3.2 Inner hair cells had normal numbers of afferent synapses, but enlarged presynaptic ribbons after noise exposure

IHCs contain presynaptic ribbons that appose postsynaptic glutamate receptors on Type I afferent auditory nerve fibers. In Figure 3, IHC presynaptic puncta are labeled with CtBP2 (red) and postsynaptic glutamate receptor puncta are labeled with GluA2 (green). Synapse counts per IHC are plotted as a function of cochlear frequency place in Figure 3D-E. As reported previously (Valero et al. 2017), control synapse counts were greatest in the mid-frequencies. On average, there was no significant difference in IHC synapse counts for the 2 month or 10 month post-exposure groups compared to controls (Figure 3D and E, Table 1). However, some individual ears showed counts > 1 standard deviation below control means for frequencies near the exposure band (Figure 3D’ and E’; 3 of 10 ears at 2 months post-exposure; 7 of 20 ears at 10 months post-exposure). In general, ears showing significant synapse loss were not the same ears that showed significant OHC loss (except two ears from the long-surviving exposure group: M114R, M119R). Few orphan ribbons (i.e. without an apposing postsynaptic component) were observed (<3% orphans across 95% of the images examined, no trends by exposure group or frequency).

**Figure 3.**
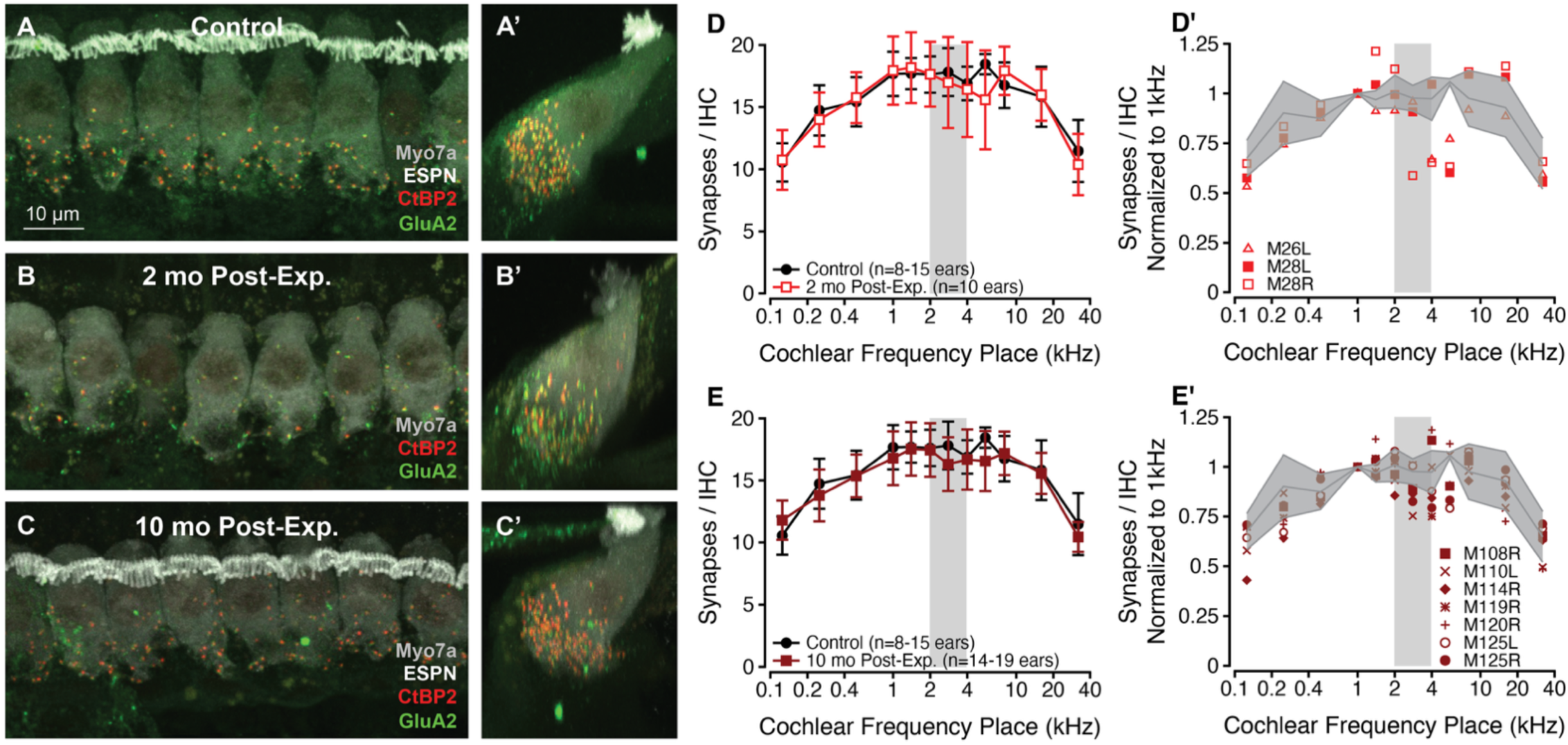
**A-C:** Confocal maximum intensity projection (MIP) images of inner hair cells at the 5.6 kHz place in a control and in subjects 2 months and 10 months post-exposure. **A,B,C**: xy MIPs. **A’,B’,C’**: yz MIPs. Note that the 2-month post-exposure (**B,B’**) case lacks espin immunostaining for stereocilia bundles. **D-E:** Mean ribbon synapses per inner hair cell as a function of cochlear frequency place in controls (black circles, *n* = 8-15 ears per frequency) and at 2 months (red open squares, *n* = 4 ears) and 10 months (dark red squares, *n* = 8 ears) post-exposure. Error bars indicate ± 1 standard deviation from the mean. **D’-E’:** Synapse counts at 2 months (**D’**) and 10 months post-exposure (**E’**), normalized within-subject to the count at the 1 kHz place. Some ears showed values > 1 standard deviation (shading) below control means (solid lines). Gray boxes illustrate the noise exposure band.

**Table 1.**
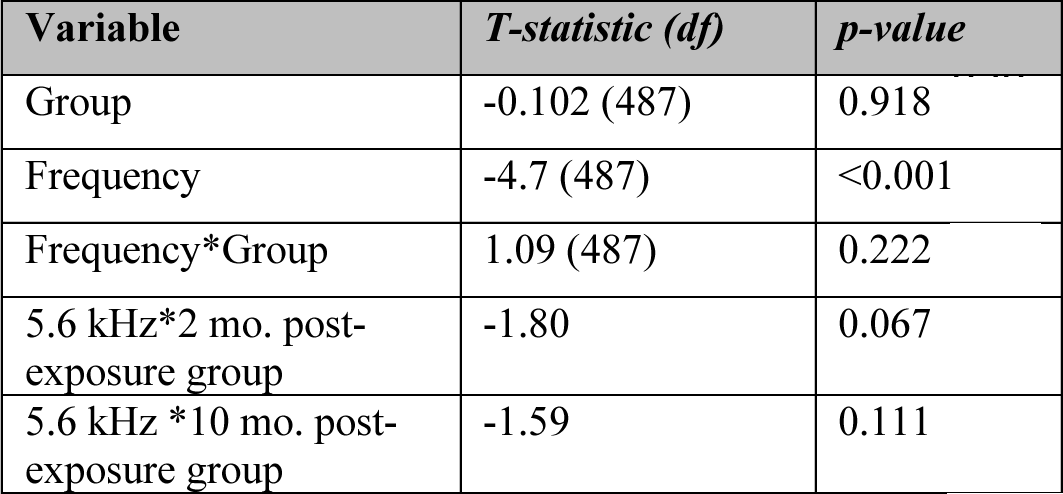
Linear mixed effects model results for IHC ribbon counts across controls, 2 months post-exposure, and 10 months post-exposure groups. *IHC Ribbon Count ∼ Group + Frequency + Group*Frequency + (1|Subject); R^2^* = 0.54

IHC ribbon volumes were measured and grouped into two-octave regions (Figure 4A). Mean volumes were similar across frequency regions and groups, however, the range increased post-exposure (Figure 4B). A jack-knife procedure was used to quantify the variability in range within each group, and to ensure the range was not driven by a few outliers (see Methods). Cumulative distribution functions were used to better illustrate this trend, and revealed clear differences in IHC ribbon volume distributions across groups (Figure 4C). At 2 months and 10 months post-exposure, there was a greater proportion of enlarged ribbons compared to controls, as shown by the rightward shift and long upper tail in the cumulative distribution functions (red lines in Figure 4C and 4C’; Table 2). These effects were present across all frequency regions (*Kolmogorov-Smirnov* tests; Table 2).

**Figure 4.**
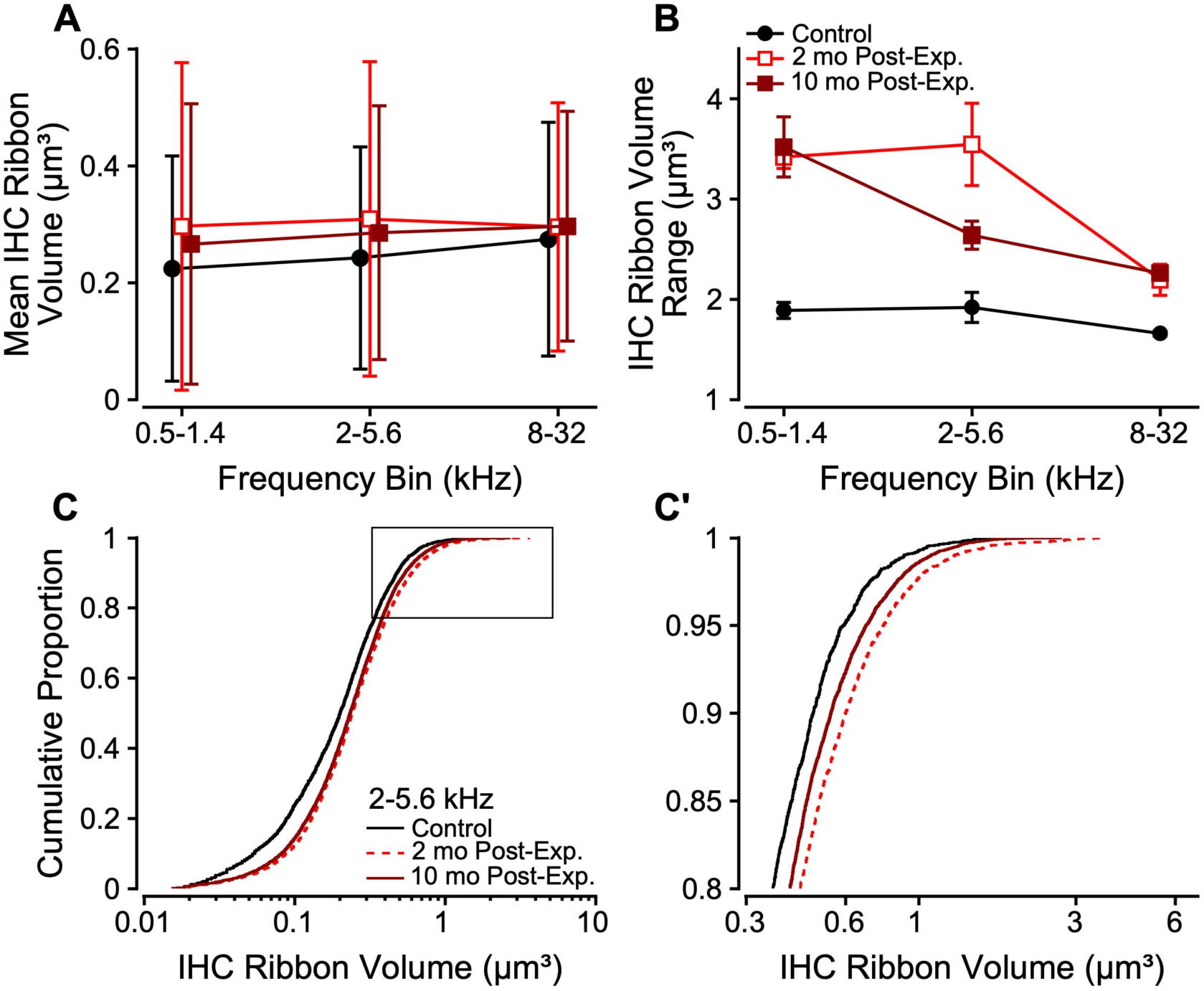
Mean (**A**) and range (**B**) of inner hair cell ribbon volume as a function of cochlear frequency in controls (black circles, *n* = 9 ears) and subjects at 2 months (open red squares, *n* = 10 ears) and 10 months (dark red squares, *n* = 19 ears) post-exposure. Error bars indicate ± 1 standard deviation from the mean. **C:** Cumulative distribution functions of IHC ribbon volume at 2-5.6 kHz in controls (black) and at 2 months (red dashed), and 10 months post-exposure (dark red). The upper tail of the distribution (box) is enlarged in **C’**.

**Table 2.** Kolmogorov-Smirnov tests comparing IHC ribbon volume distributions from controls, 2 months post-exposure, and 10 months post-exposure groups.

| Frequency (kHz) | 2 Mo. Post-Exp. |  | 10 Mo. Post-Exp. |  |
| --- | --- | --- | --- | --- |
|  | K-statistic | p-value | K-statistic | p-value |
| 1 | 0.119 | 2*10<sup>-13</sup> * | 0.189 | 1*10<sup>-39</sup> * 891 |
| 2 | 0.170 | 3*10<sup>-27</sup> * | 0.201 | 1*10<sup>-45</sup> * 892 |
| 4 | 0.277 | 3*10<sup>-64</sup> * | 0.212 | 1*10<sup>-47</sup> * 893 |
| 5.6 | 0.158 | 3*10<sup>-9</sup> * | 0.135 | 1*10<sup>-11</sup> * 894 |
| 8 | 0.195 | 2*10<sup>-32</sup> * | 0.213 | 3*10<sup>-47</sup> * 895 |
| 32 | 0.115 | 8*10<sup>-7</sup> | 0.128 | 2*10<sup>-9</sup> * 896 |

\*significant after Benjamini-Hochberg correction

### 3.3 Outer hair cell ribbons were also enlarged after noise exposure

OHCs also have afferent synapses comprised of presynaptic ribbons that appose Type II auditory nerve fibers. In Figure 2, presynaptic puncta in OHCs are also labeled by CtBP2 (red). There are fewer ribbons per OHC compared to IHCs and the number of ribbons per OHC decreased from the low-frequency to the high-frequency regions of the cochlea (Figure 5A). Compared to controls, OHC ribbon counts were significantly reduced at 2 months post-exposure, but not at 10 months post-exposure (Table 3). It should be noted that the control group was limited (*n =* 3) for this analysis.

**Figure 5.**
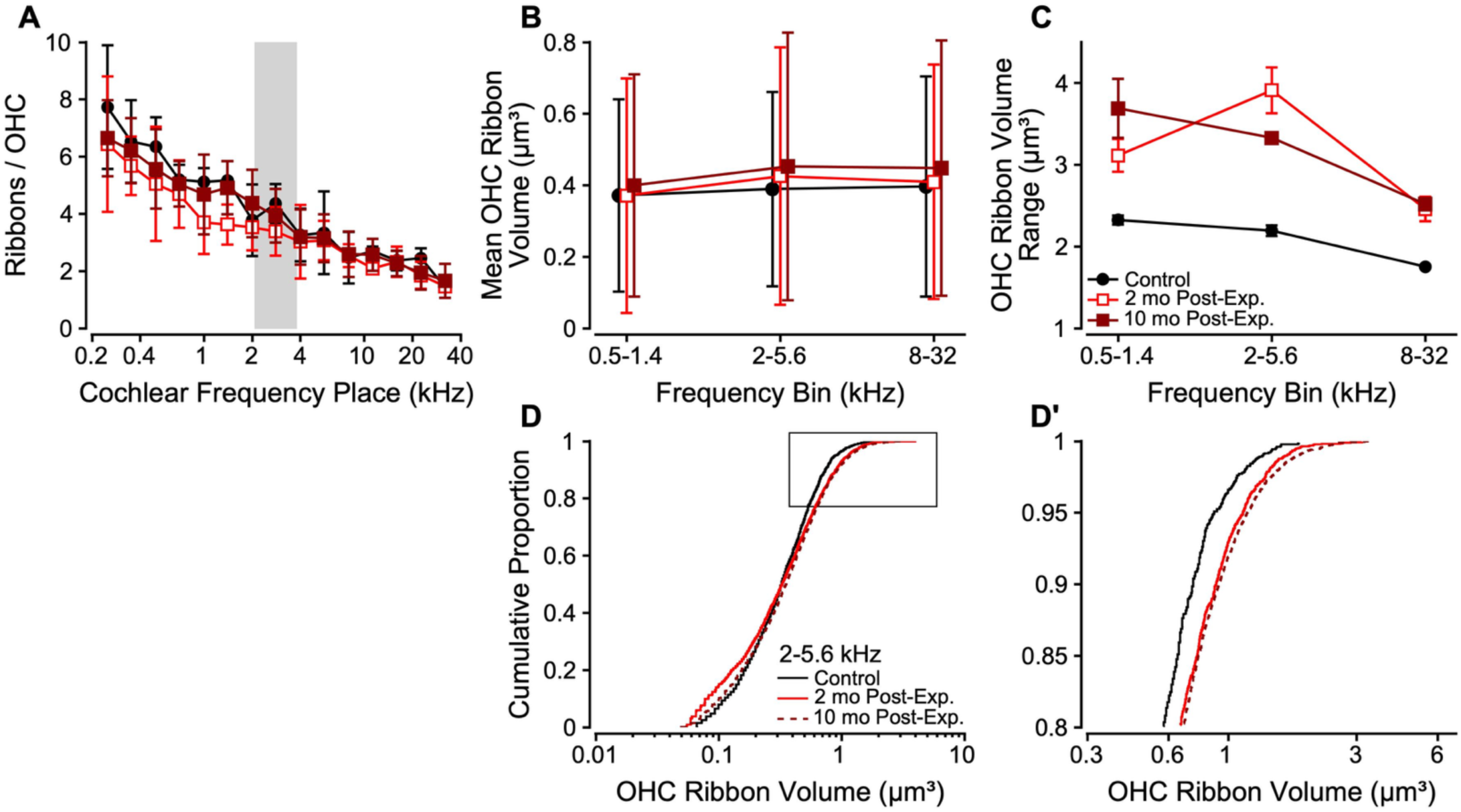
**A:** Mean ribbons per outer hair cell as a function of cochlear frequency place in controls (black circles, *n* = 3 ears) and at 2 months (open red boxes, *n* = 6) and 10 months (dark red squares, *n* = 18 ears) post-exposure. **B-E**: Mean (**B**) and range (**C**) of OHC ribbon volumes as a function of cochlear frequency region. Error bars indicate ± 1 standard deviation from the mean. **D:** Cumulative distribution functions of OHC ribbon volume at 2-5.6 kHz in controls (black) and at 2 months (red dashed) and 10 months post-exposure (dark red). The upper tail of the distribution (box) is enlarged in **D’**.

**Table 3.** Linear mixed effects model for OHC ribbon counts across controls, 2 months post-exposure, and 10 months post-exposure groups. *OHC Ribbon Count ∼ Group + Frequency + Group*Frequency + (1|Subject); R^2^* = 0.71

| Variable | *T*-statistic (df) | *p*-value |
| --- | --- | --- |
| Frequency | -5.66 (75) | < 0.001 905 |
| 2 mo. post-exposure group | -3.7 (75) | < 0.001 906 |
| 10 mo. post-exposure group | -1.46 (75) | 0.148 907 |

Volumes were measured for all ribbons across OHC rows at a given cochlear frequency place (Figure 5B). As for IHCs, OHC ribbon volumes were similar across frequencies and groups, and highly variable. However, the range of OHC ribbon volumes was larger at 2 months and 10 months post-exposure compared to controls (Figure 5C). Cumulative distribution functions revealed significant differences in OHC ribbon volume distributions across groups (Figure 5D). At 2 and 10 months post-exposure, there was a greater proportion of larger ribbons and an increased range in ribbon volume compared to controls across frequency regions, as shown by the rightward shift and flattening of the cumulative distribution functions (Figure 5D and D’, Table 4).

**Table 4.** Kolmogorov-Smirnov tests comparing OHC ribbon volume distributions from controls, 2 months post-exposure, and 10 months post-exposure groups.

| Frequency<br>(kHz) | 2 Mo. Post-Exp. |  | 10 Mo. Post-Exp. |  |
| --- | --- | --- | --- | --- |
|  | *K*-statistic | *p*-value | *K*-statistic | *p*-value |
| 1 | 0.106 | $1 \times 10^{-5} *$ | 0.064 | $0.0038 * 915$ |
| 2 | 0.084 | $0.0052 *$ | 0.092 | $1 \times 10^{-4} * 916$ |
| 4 | 0.073 | 0.052 | 0.087 | $0.0018 * 917$ |
| 5.6 | 0.164 | $1 \times 10^{-8} *$ | 0.160 | $7 \times 10^{-11} * 918$ |
| 8 | 0.124 | $8 \times 10^{-4} *$ | 0.115 | $3 \times 10^{-4} * 919$ |
| 32 | 0.126 | $0.019 *$ | 0.206 | $2 \times 10^{-7} * 920$ |

\*significant after Benjamini-Hochberg correction

### 3.4 Cholinergic efferent innervation density was unchanged after noise exposure

Olivocochlear efferent neurons project from the brainstem back to the cochlea to modulate responses to incoming sound. These cholinergic neurons can be visualized by labeling with choline acetyltransferase (ChAT; Figure 6 & 7). Medial olivocochlear (MOC) neurons innervate the OHCs, whereas lateral olivocochlear (LOC) neurons innervate Type I afferent auditory nerve fibers near their synapse with the IHCs (Guinan, 2006). MOC and LOC innervation densities were estimated from the area of ChAT staining within the OHC or IHC region, respectively (Figures 6 & 7; after Liberman & Liberman, 2019).

**Figure 6.**
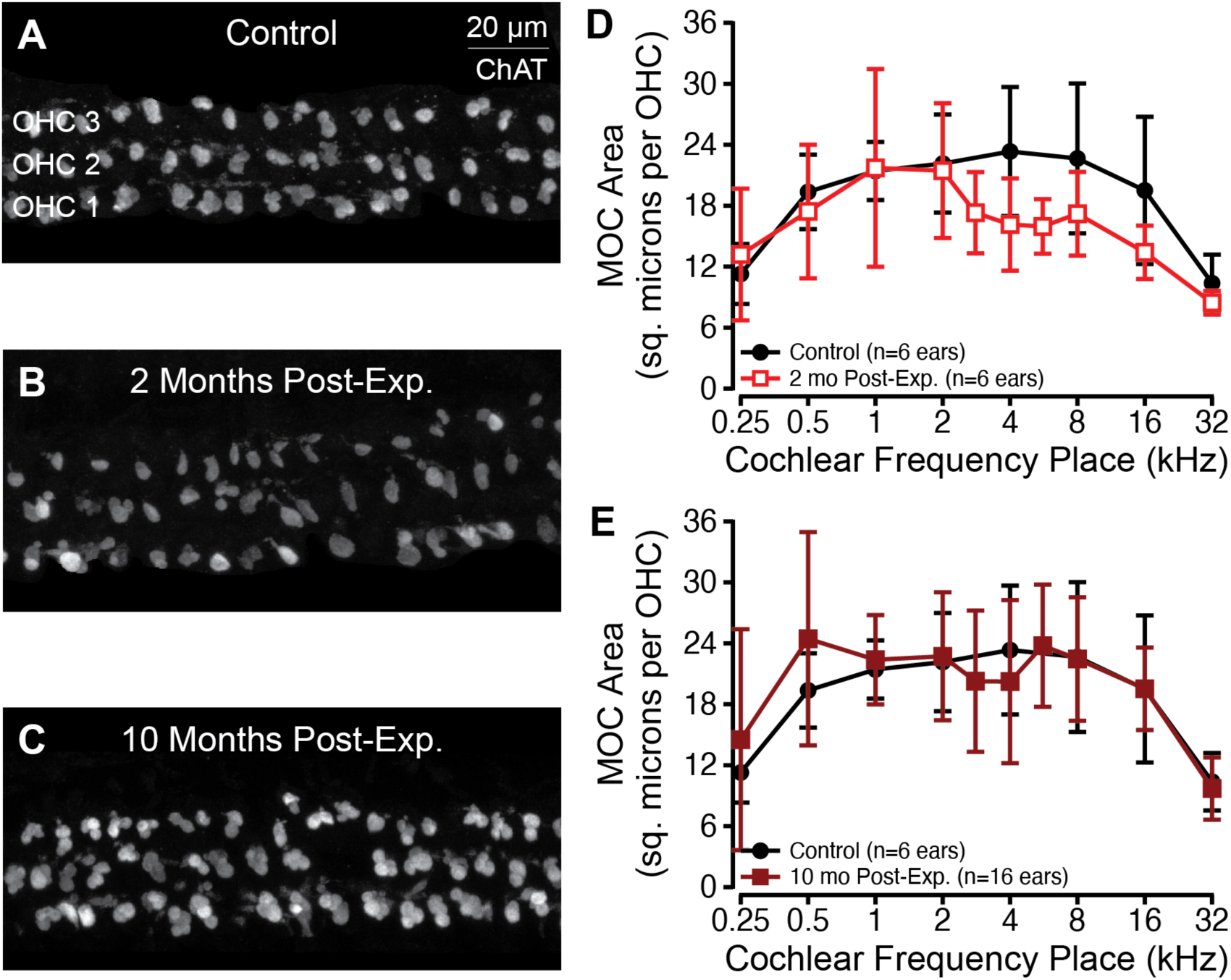
**A-C:** Confocal maximum intensity projection images of medial olivocochlear terminals labeled with ChAT at the 5.6 kHz place in a control (**A**) and exposed ear at 2 months (**B**) or 10 months (**C**) post-exposure. **D-E:** Mean medial olivocochlear terminal area (square microns per OHC) as a function of cochlear frequency for controls (black circles, *n* = 6) and subjects at 2 months (**D**; open red squares, *n* = 6 ears) and 10 months post-exposure (**E**; dark red squares, *n* = 16 ears) months post-exposure. Error bars indicate ± 1 standard deviation from the mean.

MOC innervation was densest in the mid- to high-frequency regions (Figure 6D & E, black), as previously reported in humans, cats, and rodents (Liberman & Liberman, 2019; Liberman et al., 1990). At 2 months post-exposure, MOC innervation appeared reduced at frequency places near the noise exposure band, but this was not statistically significant (Figure 6D, open red; *t*(*df*) = -1.13(106), *p* = 0.26). By 10 months post-exposure, this reduction was no longer apparent and there were no significant effects of noise exposure on MOC innervation density (Figure 6E, dark red; *t*(*df*) = 1.12(202), *p* = 0.27).

LOC innervation density decreased from apex to base (Figure 7D & E), as previously reported in humans and cats (Liberman & Liberman, 2019; Liberman et al., 1990). After noise exposure, mean LOC innervation densities were lower for some mid-frequencies. However, these differences were not statistically significant at 2 months (Figure 7D, open red; *t*(*df*) = -1.31(99), *p* = 0.19) or 10 months post-exposure (Figure 7E, dark red; *t*(*df*) = -1.68(171), *p* = 0.09).

**Figure 7.**
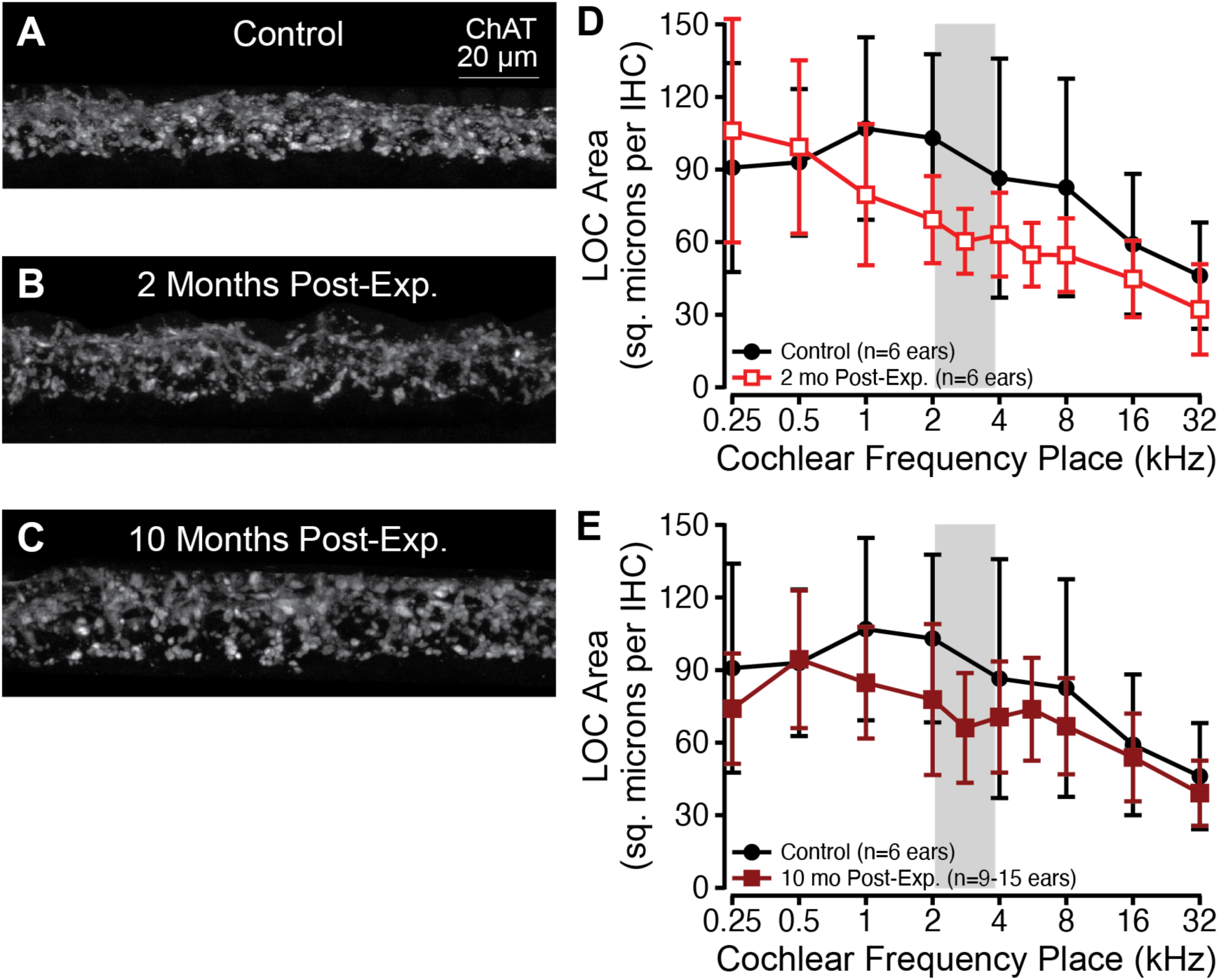
**A-C:** Confocal maximum intensity projection images of lateral olivocochlear terminals labeled with ChAT at the 5.6 kHz place in a control (**A**) and exposed ear at 2 months (B) or 10 months (**C**) post-exposure. **D-E:** Mean lateral olivocochlear terminal area (square microns per IHC) as a function of cochlear frequency for controls (black circles, *n* = 6) and subjects at 2 months (**D**; open red squares, *n* = 6 ears) and 10 months post-exposure (**E**; dark red squares, *n* = 16 ears) months post-exposure. Error bars indicate ± 1 standard deviation from the mean.

## 4. DISCUSSION

Here, we described cochlear histology in rhesus macaques with and without acoustic overexposure that caused temporary threshold shifts. First, we contributed a detailed view of the normal macaque inner ear to the literature at a time when increasing emphasis is placed on translational animal models in biomedical research, including auditory research (Capshaw et al., 2023). The macaque cochlea exhibits many similarities to the human, which highlights the utility of nonhuman primates in studies of hearing and auditory pathologies (Burton et al., 2019). Second, we established a macaque model of noise-induced SYN that is defined by chronic enlargement of both IHC and OHC ribbons and transient loss of OHC ribbons, in the absence of significant IHC and OHC loss, IHC ribbon synapse loss, or changes to cholinergic MOC and LOC innervation density. In addition, as shown in previous studies of noise-induced hearing loss in nonhuman primates by our group (Burton et al., 2020; Hauser et al., 2018; Mackey et al., 2021; Valero et al., 2017) and others (reviewed in Burton et al., 2019), macaques exhibit a high degree of inter-subject variability in their response to noise exposure.

### 4.1 Normal cochlear anatomy in the rhesus macaque

Macaques have three to four rows of OHCs and one row of IHCs (Figure 2; Johnsson & Hawkins, 1967; Lonsbury-Martin et al., 1988) like humans (Bredberg, 1968; Johnsson & Hawkins, 1967). Macaque IHC ribbon synapse counts were similar to rodents (Furman et al., 2013; Hickox et al., 2017; Liberman & Liberman, 2015, 2016) and estimates in humans (Wu et al., 2019), showing the characteristic inverted-U shape as a function of frequency. Macaque OHC ribbon counts decreased with increasing frequency as in cats (Liberman et al., 1990), whereas OHC ribbon counts in mice are similar across frequencies (Liberman & Liberman, 2016; Wood et al., 2021). The functional significance of this species difference is unknown, due to limited knowledge of the role of OHC ribbons and their apposing Type II afferent fibers.

Macaque IHC and OHC ribbons had variable volumes, as previously reported in other species (Weisz et al., 2012). The relative size of IHC and OHC ribbons seems to vary across species. Cats have larger IHC ribbons than OHC ribbons (Liberman et al., 1990); rats have similar IHC and OHC ribbon volumes (Weisz et al., 2012); and macaques have significantly smaller IHC ribbons than OHC ribbons (*k* = 0.34, *p* < 0.001). Whether these differences in OHC vs. IHC synaptic body size and shape translate to functional differences is unknown.

In macaques, MOC innervation density was greatest in the mid-frequencies, consistent with most other animals and humans (Liberman & Liberman, 2019; Steenken et al., 2024). Macaques have intermediate MOC innervation density compared to that of rodents (dense) vs. humans (sparse). Macaque LOC innervation density was greatest for low- to mid-frequencies, similar to gerbils (Steenken et al., 2024) and humans (Liberman & Liberman, 2019).

### 4.2 Cochlear histopathology following noise: Species comparisons and effects of post-exposure survival

#### 4.2.1 Outer and inner hair cell survival

Rodent models of noise-induced SYN secondary to temporary threshold shifts exhibit IHC ribbon synapse loss in the absence of hair cell loss (e.g. Kujawa & Liberman, 2009). In the present study, macaques also had minimal OHC and IHC loss at 2 and 10 months after temporary noise-induced threshold shifts. A few macaque ears exhibited substantial OHC loss, consistent with higher inter-animal variability in the response to acoustic injury that is common to genetically heterogeneous species, such as guinea pigs and macaques, compared to inbred mice (Wang et al., 2002).

Hook lesions, or complete hair cell loss in the basal tip of the cochlea, are reported in some models of noise-induced SYN (Liberman & Liberman, 2015). There was no evidence of a hook lesion in our macaques, as there was only modest OHC and IHC loss in the basal-most cochlear frequency region.

#### 4.2.2 Inner hair cell ribbon synapse counts

IHC ribbon synapse loss occurs secondary to glutamate excitotoxicity (Hu et al., 2020; Kim et al., 2019). Although some individual ears showed significant IHC synapse loss, our noise exposure did not significantly affect group-mean synapse counts at 2 months or 10 months post-exposure. Furthermore, even in the few maximally affected ears, IHC ribbon synapse loss only reached 45% at 2 months post-exposure and 25% at 10 months post-exposure, which is still less than the ∼50% synapse loss typically observed in rodent models (Bharadwaj et al., 2021; Kujawa & Liberman, 2009; Lin et al., 2011; Singer et al., 2013). However, given the high cost of nonhuman primates (Burton et al., 2019), it was not feasible to titrate noise level in the same way possible with rodents (Fernandez et al., 2020). Thus, in contrast to most prior studies that produced noise-induced SYN (e.g., Kujawa & Liberman, 2009; Lin et al., 2011), where exposure level was set at the maximum possible (for the given spectrum and duration) without producing PTS, we do not know how close our noise exposure was to this critical level. In rodents, it is clear that there are many TTS-producing noise exposures that do not cause SYN (Fernandez et al., 2020).

The lack of significant IHC synapse loss may also be a product of the relatively long post-exposure survival times examined in this study. Synapse counts recover over time following acoustic overexposure in guinea pigs (Hickman et al., 2020, 2021; Shi et al., 2013; Song et al., 2016) and some mouse strains (Kim et al., 2019; Kujawa & Liberman, 2009; Liberman & Liberman, 2015; Shi et al., 2015; Wu et al., 2024). Much of this synaptic repair/regeneration can occur within 1 month post-exposure, and our 2 month and 10 month post-exposure time points exceed this window. Therefore, it is possible that our macaques experienced immediate synapse loss followed by partial to full recovery by the time of histological analysis, although there is no direct evidence for this interpretation.

Despite the absence of IHC ribbon synapse loss, we did observe significant, permanent changes in physiological function and perceptual abilities in our noise-exposed macaques (Conner et al., 2025; Mackey et al., 2025). These data suggest that even a mild or potentially temporary loss of synapses accompanied by synaptic hypertrophy can have lasting effects on auditory physiology and perception. Repaired or regenerated synaptic physiology may differ from innate/native synapses (Vincent et al., 2022), contributing to changes in central auditory function and hearing deficits. Furthermore, SYN-related changes to central auditory physiology, such as the inferior colliculus (Bakay et al., 2018; Bakay et al., 2024; Shaheen & Liberman, 2018) and auditory cortex (Asokan et al., 2018; Resnik & Polley, 2021), may not be reversed by peripheral repair.

#### 4.2.3 Inner hair cell ribbon volumes

Enlarged IHC ribbon volumes have been observed in noise-induced SYN and age-related hearing loss (Blum et al., 2024; Furman et al., 2013; Hickman et al., 2020; Jeng, Ceriani, et al., 2020; Kim et al., 2019; Song et al., 2016; Stamataki et al., 2006; Valero et al., 2017). Ribbon enlargement occurs immediately following noise exposure (Liberman & Liberman, 2015), but is independent of glutamate excitotoxic processes (Kim et al., 2019). Sustained ribbon enlargement is consistently observed up to 1-4 weeks post-exposure (Blum et al., 2024; Furman et al., 2013; Hickman et al., 2020; Kim et al., 2019; Liberman & Liberman, 2015; Song et al., 2016), may be evident in guinea pigs through 1-6 months post-exposure (Hickman et al., 2020; Song et al., 2016), at 2 months post-exposure in macaques (Valero et al., 2017), and now appears as a persistent phenotype through even later post-exposure times (up to 10 months) in macaques.

The functional significance of enlarged ribbons is unknown, but likely affects synaptic physiology and auditory nerve function. The capacity for tethering and fusing readily releasable vesicles varies with ribbon size (Becker et al., 2018; Matthews & Fuchs, 2010; Moser et al., 2020). Larger ribbons may tether more vesicles (Sheets et al., 2017; Song et al., 2016), yielding greater multivesicular release and larger postsynaptic potentials, leading to enhanced onset response in auditory nerve fibers and therefore the larger Wave I amplitudes observed in this study (Figure 1C). The larger CtBP2 puncta seen in the confocal images may actually represent the presence of multiple pre-synaptic ribbons at a single synapse, as is frequently observed at the ultrastructural level in the synaptopathic mouse (Moverman et al., 2023).

While the classical presentation of noise-induced SYN is accompanied by reduced ABR Wave I amplitudes (Kujawa & Liberman, 2009) and mixed effects on single-unit auditory nerve responses (Furman et al., 2013; Song et al., 2016), these findings are for stimuli in quiet, at shorter post-exposure times when synapse loss is still present, or in mice that show permanent synapse loss. In two recent studies by Suthakar and Liberman (2021, 2022), synaptopathic mice showed enhanced neural responses to tones in noise, via single-unit auditory nerve recordings and ABR. The SYN model used in their studies also shows enlarged IHC ribbon volumes (Liberman & Liberman, 2015), supporting the hypothesis that enlarged or multiplexed ribbons facilitate increased sound-evoked auditory nerve activity (Jeng, Ceriani, et al., 2020).

#### 4.2.4 Outer hair cell ribbon synapses

Relatively few studies have focused on OHC ribbons, especially in the context of noise exposure. The OHC to Type II auditory-nerve fiber synapse does not appear to experience glutamate excitotoxicity as does the IHC afferent synapse. Type II fibers do not express the same type of AMPA receptors as Type I fibers (Liberman et al., 2011) and do not exhibit terminal swelling under conditions that are excitotoxic to Type I afferents.

Wood et al. (2021) provided the first characterization of OHC ribbons in a mouse model of noise-induced SYN. At 7 days post-exposure, mice exhibited increased counts and sizes of OHC ribbons compared to controls. Temporary threshold shifts had not yet resolved, and OHC loss was not significantly different from controls. In contrast, OHC ribbon counts were reduced in a mouse model of age-related hearing loss, but ribbon size was not characterized (Jeng, Johnson, et al., 2020).

Our macaques showed a transient reduction in OHC ribbon counts at 2 months post-exposure, and persistently enlarged OHC ribbon volumes at 2 and 10 months post-exposure. Perhaps the increase in OHC ribbon count in the Wood et al. study is an acute response to acoustic injury, whereas enlarged OHC ribbons may reflect a chronic change. As with IHC ribbons, the functional significance of enlarged OHC ribbons is not known. Type II afferent signaling can be impacted by noise exposure (Nowak et al., 2021), though the functional contribution of these neurons, including a role in indicating cochlear damage, is debated (Liu et al., 2015; Maison et al., 2016; Weisz et al., 2012; Zhang & Coate, 2017).

#### 4.2.5 Efferent terminal density

Olivocochlear efferent neurons project from the brainstem to the cochlea and modulate cochlear responses to incoming sound (Guinan, 2018). The MOC system reduces OHC motility to enhance responses to acoustic transients in the presence of continuous masking noise (Guinan, 2006; Lopez-Poveda, 2018; Wiederhold & Kiang, 1970; Winslow & Sachs, 1987). Sound-evoked MOC feedback also reduces damage from acoustic overexposure by mechanisms that are less clear (Maison et al., 2007). MOC neurons are thought to receive input from low spontaneous rate ANFs (Liberman, 1988), which may be preferentially lost in SYN (Furman et al., 2013; Liberman & Liberman, 2015; Schmiedt et al., 1996; but see Suthakar & Liberman, 2021). Thus, MOC function could be impacted by IHC synapse loss. Some studies in rodents show reduced MOC innervation density following noise-induced SYN (Boero et al., 2018; Qian et al., 2021; Steenken et al., 2024) or age-related hearing loss (Boero et al., 2020; Grierson et al., 2022; Jeng, Johnson, et al., 2020), and reduced MOC innervation in aging humans with minimal OHC loss (Liberman & Liberman, 2019). However, others find no change in MOC innervation density in noise-exposed (Grierson et al., 2022) or aging mice (Kobrina et al., 2020). In the present study, we saw no significant changes to MOC innervation at 2 months or 10 months post-exposure. Etiology and time course of inner ear damage may differentially affect the MOC system.

Less is known about the unmyelinated LOC neurons, which synapse onto the dendrites of Type I auditory nerve fibers (Guinan, 2018). As seen in the present data, cholinergic LOC innervation density appears minimally affected by noise (Grierson et al., 2022) and age (Kobrina et al., 2020; Liberman & Liberman, 2019; Steenken et al., 2024), although the location of LOC terminals may shift from afferent dendrites to IHCs in aging (Jeng et al., 2021; Lauer et al., 2012). LOC neurotransmitter expression can be altered by noise exposure, such as through the upregulation of dopamine precursors (Frank et al., 2023; Niu & Canlon, 2002; Wu et al., 2020, but see Grierson et al., 2022), pointing to other possible forms of noise-induced plasticity.

### 4.3 Implications for diagnostic testing

Anatomical characterization of SYN identifies sites of lesion to guide the development of diagnostic testing and therapeutic treatment strategies. Unlike SNHL, which can be readily diagnosed using otoacoustic emissions assays of OHC function, the more subtle cochlear changes accompanying SYN remain hidden from current clinical diagnostic approaches. If enlarged and/or multiplexed synaptic ribbons are indeed a prominent and permanent consequence of SYN, then physiological or psychophysical measures that probe concomitant changes in synaptic physiology would be promising as biomarkers for SYN.

## ACKNOWLEDGEMENTS

The authors wish to acknowledge Mary Feurtado for her assistance with anesthesia administration and monitoring, as well as Alejandro Tarabillo and Jessica Feller for assisting with physiological data collection. This work was supported by the National Institutes of Health (R01 DC 015988 (RR), NIH F32 DC 019817 (JAM), NIH F31 DC 019823 (CAM)) and Fondation pour l’Audition (MCL).

